# Transcranial alternating current stimulation induces long-term augmentation of neural connectivity and sustained anxiety reduction

**DOI:** 10.1101/204222

**Authors:** Kevin J. Clancy, Sarah K. Baisley, Alejandro Albizu, Nicholas Kartvelishvili, Mingzhou Ding, Wen Li

## Abstract

Although transcranial alternating current stimulation (tACS) has demonstrated short-term effects in modulating neural oscillations, potential clinical efficacy of tACS for treating “oscillopathies” (disorders associated with aberrant neural oscillations) hinges on its ability to generate long-term neural plasticity, which can translate into lasting behavioral changes. Administering alpha-frequency tACS over 4 consecutive days, we evaluated short- and long-term (> 24 hours) effects of α-tACS on local alpha power and oscillatory connectivity, along with behavioral outcomes in anxious arousal and affective sensory perception. The α-tACS (vs. sham stimulation) group exhibited increases in posterior alpha power immediately and 30 minutes post-stimulation but not 24 hours post-stimulation, suggesting transient effects on local neuronal synchrony. Strikingly, long-range alpha-frequency Granger causal connectivity (posterior→frontal) increased not only immediately and 30 minutes but also 24 hours post-stimulation, paralleled by sustained reductions in anxious arousal and aversion to auditory stimuli. Therefore, tACS is capable of eliciting long-term plasticity in long-range oscillatory connectivity with direct, lasting behavioral consequences, while leaving endogenous local oscillations unaltered. This temporal disparity in local and network effects favors the view of large-scale network strengthening by tACS, calling for research attention to the circuit and network impacts of tACS. Critically, with the growing recognition of large-scale network dysfunctions as a transdiagnostic pathophysiology of psychiatric disorders, this connectivity plasticity advocates for the clinical application of tACS with its unique advantage in tackling network pathologies.

## SIGNIFICANCE STATEMENT

Aberrant neural oscillations have been increasingly identified in a variety of neuropsychiatric disorders (also known as “oscillopathies”). Consistent demonstration of modulated neural oscillations via transcranial alternating current stimulation (tACS) has raised the enticing possibility of applying this new technology as a novel, non-invasive treatment for oscillopathic disorders. Using a multi-session alpha-frequency tACS protocol, we provide evidence for longterm effects of tACS in augmenting cortico-cortical oscillatory connectivity, which is further accompanied by sustained reductions in anxious arousal and aversion to sensory stimuli. Such lasting changes in both neural and behavioral responses underscore the clinical feasibility and efficacy of tACS, especially in treating conditions rooted in dysfunctional neural networks, a transdiagnostic pathophysiology in multiple mental disorders.

## INTRODUCTION

Brain oscillations play important roles in supporting various mental activities (e.g., working memory, attention, sensory processing) by regulating neuronal excitability and maintaining neural communication over local circuits and global networks (1–9). Aberrations in such oscillatory activity have been observed in a host of neuropsychiatric disorders (10, 11), potentially underpinning their neuropathophysiology. Recent advances in transcranial alternating current stimulation (tACS) demonstrate that by applying weak electric fields tuned to the rhythms of endogenous oscillations through the scalp, tACS can directly interact with and thereby modify brain oscillations (12–15). This non-invasive neuromodulation technology thus holds great promise for therapeutic interventions in neuropsychiatric disorders characterized by oscillopathies (16). However, clinical efficacy of tACS hinges on the premise that its effects persist and induce long-term neural plasticity and lasting behavioral changes.

To the extent that tACS has shown reliable aftereffects, i.e., increased power of the targeted oscillations, lasting fairly consistently for 30 minutes (17–19), long-term effects (> 24 hours) have not been reported. Existing evidence rather indicates that the enhanced power disappears following an extended window of observation (e.g., 70 minutes) (19). Whether tACS is capable of generating long-term effects would depend on the mechanisms by which it modulates neural oscillations. Current theories of tACS effects center on two putative mechanisms: (1) transient neural entrainment to an external driving rhythm (12–14, 20), and (2) neural plasticity, by which long-lasting enhancement of the stimulated oscillations is achieved by strengthening oscillatory circuits (15, 18, 20–22). It is possible that both mechanisms are engaged by tACS (23), whereby transient synchronized spiking via neural entrainment may facilitate spike-timing dependent plasticity (STDP) at the synapses (24), promoting long-lasting tACS effects.

Arguably, the limited durability of tACS effects to date could merely reflect a dose-effect phenomenon such that higher doses of tACS (e.g., repeated stimulation) would yield longer-lasting tACS effects. Nevertheless, if the underlying mechanism is primarily neural entrainment to exogenous oscillatory electric fields, the effects would be reversed by endogenous generating mechanisms (especially by deep generators that are largely impervious to tACS) upon cessation of exogenous stimulation. For example, tACS at the alpha frequency (i.e., α-tACS), one of the most-studied tACS techniques, mainly stimulates the occipito-parietal cortex (13, 15, 25, 26), rarely reaching deep generators in subcortical structures such as the thalamus (27). Given that the thalamo-cortical circuit operates as a unified generator of alpha oscillations (28, 29), deep thalamic driving inputs can reset cortical alpha oscillatory activity after the cessation of α-tACS.

Conversely, by the second mechanism, tACS can generate lasting neural plasticity via strengthened circuits, especially by recruiting the abundant recurrent cortico-cortical connections whose reverberations would modify large-scale network dynamics (23, 30). Particularly, α-tACS effects can transpire via the alpha oscillatory circuits that are instrumental for long-range interregional interactions (8, 31, 32). Such cortico-cortical connectivity changes are not only conducive to lasting neural plasticity but also would be relatively independent of, and thus resistant to, the rectifying inputs from deep thalamic generators. Accordingly, cortico-cortical connectivity plasticity would not only parallel but may also outlast local power enhancement.

Here, our study aimed to evaluate the long-term effects of repeated α-tACS across four consecutive days (Figure 1B) in modulating local alpha power and long-range alpha-frequency connectivity (using Granger causality analysis) (33, 34), particularly posterior→frontal connectivity that dominates resting-state alpha communication (2, 35–37). To assess the clinical outcome of tACS, we measured changes in anxious arousal, which is closely related to local and inter-areal alpha oscillatory activities (11, 38–40). Also, as auditory and olfactory processing and perception are particularly sensitive to states of arousal and alertness (41–45), we assayed changes in affective perception of simple auditory and olfactory stimuli (negative and neutral) as another clinical outcome.

**Figure 1.**
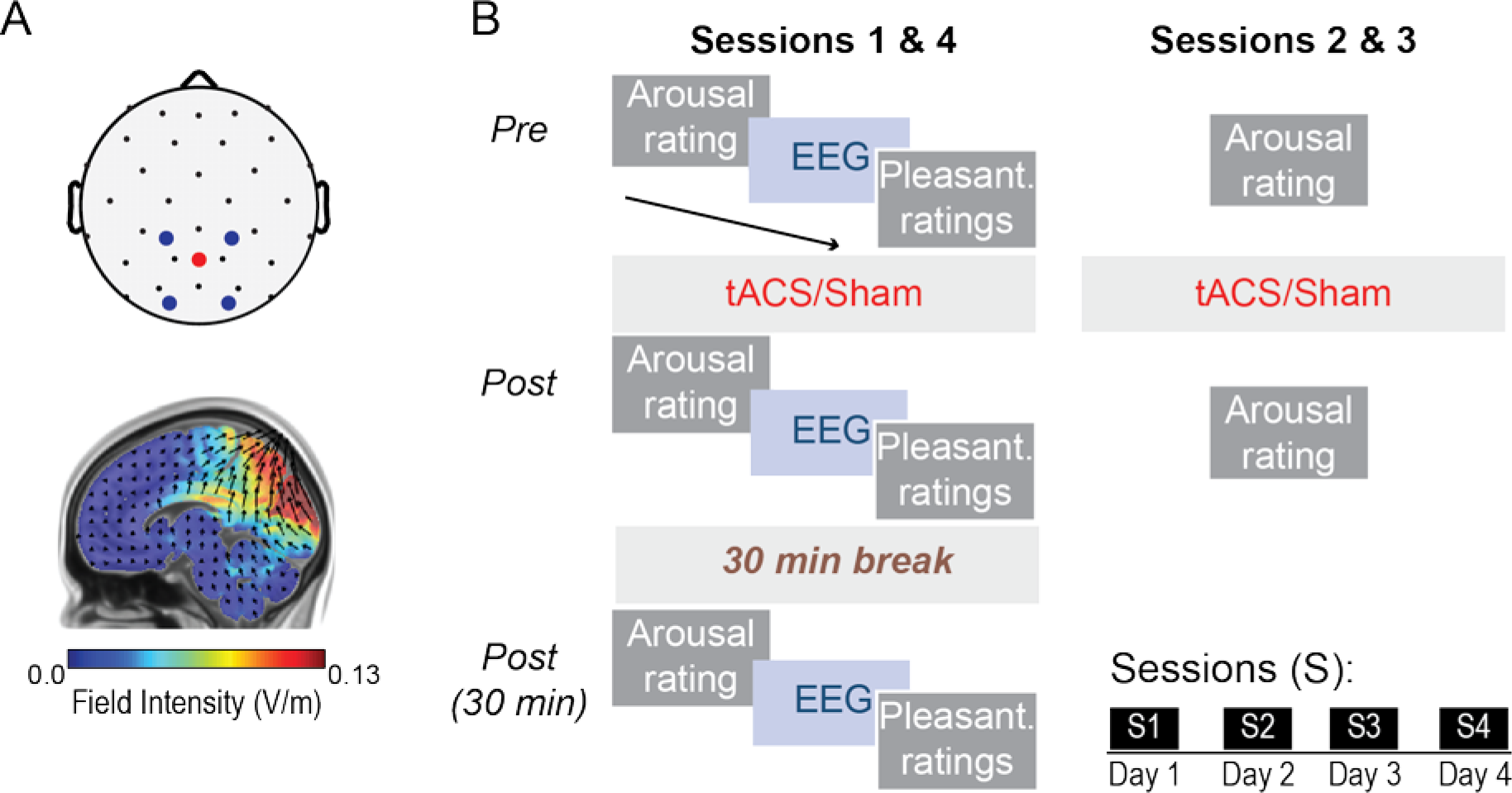
Stimulation setup and experimental protocol. A) Top: tACS montage, with stimulation electrodes placed over occipito-parietal sites. Bottom: Modeled current flow of tACS the cortex demonstrates maximum electrical field intensity over dorsal occipital cortex. B) Participants completed 4 sessions of tACS or sham stimulation, each separated by 24 hours. tACS was administered for 30 minutes at each individual’s peak alpha frequency. Eyes-open resting state EEG data and ratings of anxious arousal and perceived pleasantness of auditory and olfactory stimuli were acquired before, immediately after, and 30 minutes after stimulation.

## RESULTS

Subjective ratings of anxious arousal were acquired daily, while ratings of stimulus pleasantness and resting-state eyes-open electroencephalogram (EEG) were acquired at the first and last sessions in both the Active and Sham groups (Figure 1B). In general, we examined immediate and short-term (30 minutes) effects (in the Active vs. Sham group) based on within-session changes and assessed long-term effects based on baseline shifts between the initial and final baselines (reflecting > 24 hours from the most recent stimulation). Interactions between Sessions, Time (pre, post, post-30 minutes) and Group, i.e., how within-session effects changed from the initial to the final session (in the Active vs. Sham group), were also examined to inform the long-term effects of tACS that influenced the impact of subsequent tACS.

### tACS induced immediate and short-term increases in occipito-parietal alpha power

An omnibus repeated-measures Analysis of Variance (rANOVA) of Site (left, middle, right), Session (initial, final), Time (pre, post, post-30 minutes), and Group (active, sham) on posterior alpha power revealed a significant interaction of Time x Group (*F*_*1.99, 71.79*_ = 3.35, *p* = .041, *η*_p_^2^ = .09), such that, across occipito-parietal sites and the two sessions, an effect of Time arose in the Active group (*F*_*1.55, 30.98*_ = 7.99, *p* = .003, *η*_p_^2^ = .29) but not the Sham group (*p* = .485; Figure 2A). Follow-up contrasts within the Active group demonstrated power (collapsed across three posterior sites) increases both immediately (*p* = .0003) and 30 minutes (*p* = .006) after stimulation. However, baselines for the two sessions did not differ in either group (*p’*s > .598), failing to indicate a long-term shift in alpha power (Figure 2B). Additionally, there was no Session × Time × Group interaction (*p* = .268) such that the within-session effects did not change from the first to the last session. Overall, the results confirmed that tACS produced an immediate increase in alpha power, lasting 30 minutes post-stimulation.

**Figure 2.**
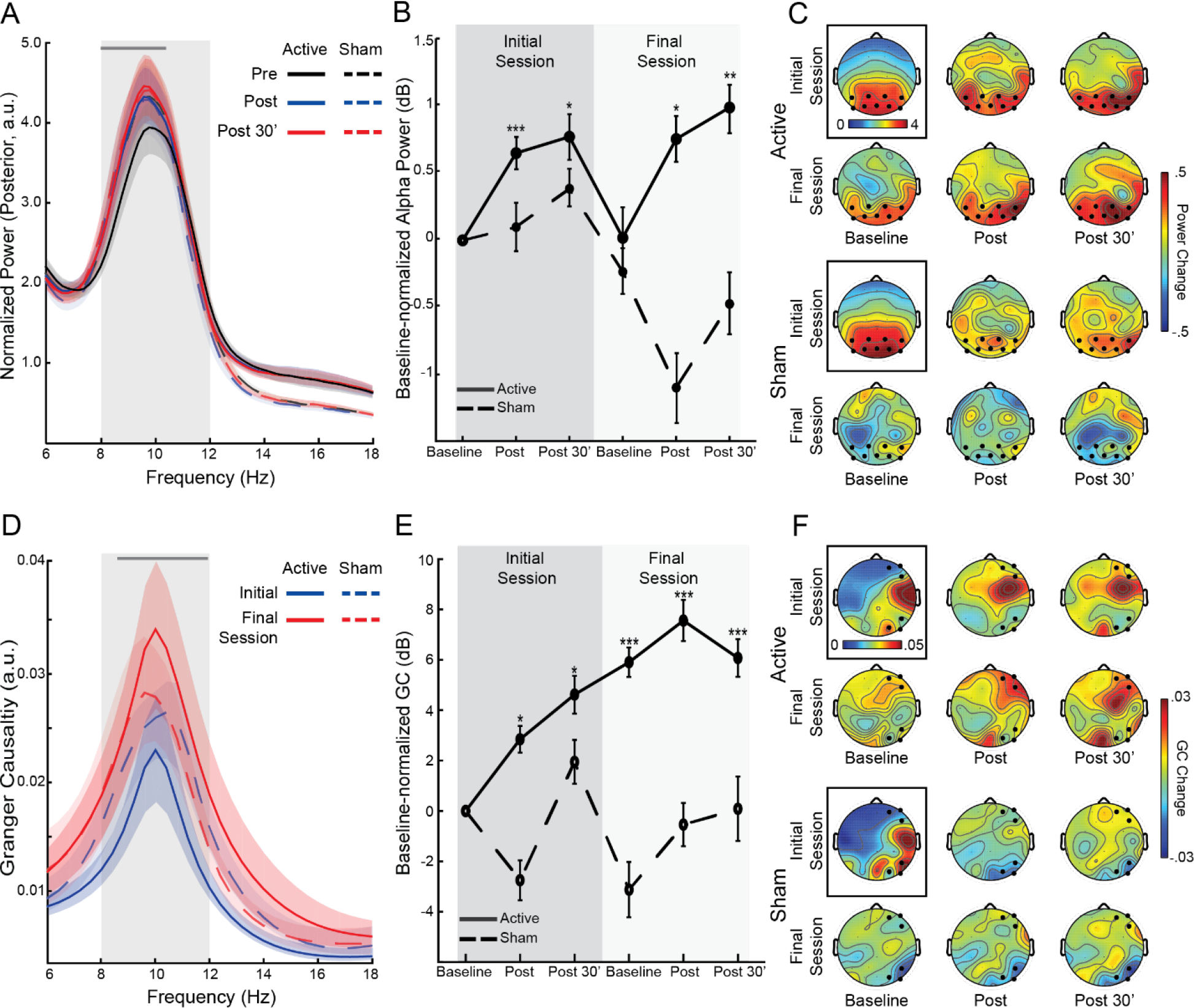
Changes in posterior alpha power and posterior→frontal alpha GC. A) Spectra of normalized power (collapsed across occipito-parietal electrodes) for each group at the three time points, averaged across the initial and final sessions. The grey line at the top of the waveforms indicates the frequency bins (.25 Hz each) showing a significant Group-by-Time interaction. The interaction effect appeared selectively in the lower half of the targeted alpha range (8-10.5 Hz). B) Alpha power magnitudes, decibel-normalized to the initial baseline, for each group. C) Scalp topographical maps of normalized alpha power at the initial baseline (surrounded by a black box and associated with its own color scale), and power changes at each assessment thereafter. Electrodes included in occipito-parietal sites are bolded. D) Spectra of right-hemisphereposterior→frontal Granger causality for each group (averaged across time for simplicity). The grey line at the top of the waveforms indicates frequency bins (.5 Hz each) showing a significant Group-by-Session interaction, which spans almost the entire alpha band (8.5-12 Hz). E) Magnitudes of right posterior→frontal alpha GC, decibel-normalized to initial baseline, for each group. F) Scalp topographical maps of alpha GC sent from right occipito-parietal electrodes at the initial baseline (surrounded by a black box and associated with its own color scale), and changes at each assessment thereafter. Sending occipito-parietal and receiving frontal electrodes are bolded. Shaded ribbons = standard error of the mean (SEM); Error bars = SEM. \**p* < .05; \*\**p* < .01; *** *p* < .005. All *p*’s are relative to the initial baseline. Note, the groups did not differ in the initial baseline values (*p’*s > 19)

### tACS induced immediate, short-term and long-term increases in posterior→frontal alpha connectivity

A similar omnibus rANOVA of Site (right, left hemisphere), Session (initial, final), Time (pre, post, post-30 minutes), and Group (active, sham) on posterior→frontal alpha GC showed a significant four-way interaction of Site × Session × Time × Group (*F*_*1.98, 71.13*_ = 4.18, *p* = .020, *η*_p_^2^ = .10). A follow-up ANOVA (Session × Time × Group) in the right hemisphere revealed a Session × Group interaction (*F*_*1, 36 =*_ 7.71, *p* = .009, *η*_p_^2^ = .18; Figure 2D), which was substantiated by a significant increase from the initial to the final session in the Active (*F*_*1, 20*_ = 12.11, *p* = .002, *η*_p_^2^ = .38) but not Sham (*p* = .487) group. Furthermore, as illustrated in Figure 2E, relative to the initial baseline, GC at all subsequent assessments was augmented in the Active group (*t*’s < −2.28, *p*’s < .05 FDR corrected) but not in the Sham group (*p*’s > .190), reflecting immediate and sustained enhancement of connectivity via tACS. Importantly, the Active group (but not the Sham group, *p* = .225) demonstrated a significant baseline shift from the initial to the final session (*F*_*1, 20*_ = 18.46, *p* = .0004, *η*_p_^2^ = .48), highlighting a long-term effect of tACS.

A similar follow-up ANOVA in the left hemisphere showed an interaction of Session × Time × Group (*F*_*1.99, 71.58*_ = 2.84, *p* = .065, *η*_p_^2^ = .07), which was substantiated by an interaction of Session × Time in the Active (*F*_*1.99, 39.81*_ = 3.13, *p* = .055, *η*_p_^2^ = .14) but not Sham (*p* = .363) group. Specifically, the Active group exhibited an effect of Time in the initial session (*F*_*1.96, 39.28*_ = 3.87, *p* = .030, *η*_p_^2^ = .16), where a trend of increased alpha GC appeared immediately after stimulation (*p* = .065) and returned to baseline 30 minutes later (*p* = .504). No effect of Time in the final session (*p* = .730) or baseline shift was observed (*p* = .367), suggesting no lasting effect of stimulation in the left hemisphere.

### tACS reduced anxious arousal

An omnibus rANOVA of Session, Time, and Group on anxious arousal ratings (acquired every day) revealed a main effect of Group (*F*_*1, 32*_ = 4.64, *p* = .039, *η*_p_^2^ = .13): following the initial baseline, the two groups diverged in anxious arousal ratings, with the Active Group showing consistently lowered ratings relative to initial baseline (*p’*s < .05 FDR corrected), while no change was observed in the Sham group (*p’*s > .317; Figure 3A). Importantly, the Active group, but not Sham (*p* = .607) demonstrated sustained baseline shifts across sessions (*F_2.60, 41.62_* = 4.76, *p* = .008, *η*_p_^2^ = .23), such that reductions in pre-stimulation anxious arousal persisted at the second (*p* = .003), third (*p* = .021), and final (*p* = .005) baselines (Figure 3A). Therefore, tACS led to a consistent and lasting reduction in anxious arousal over the 4-day period.

**Figure 3.**
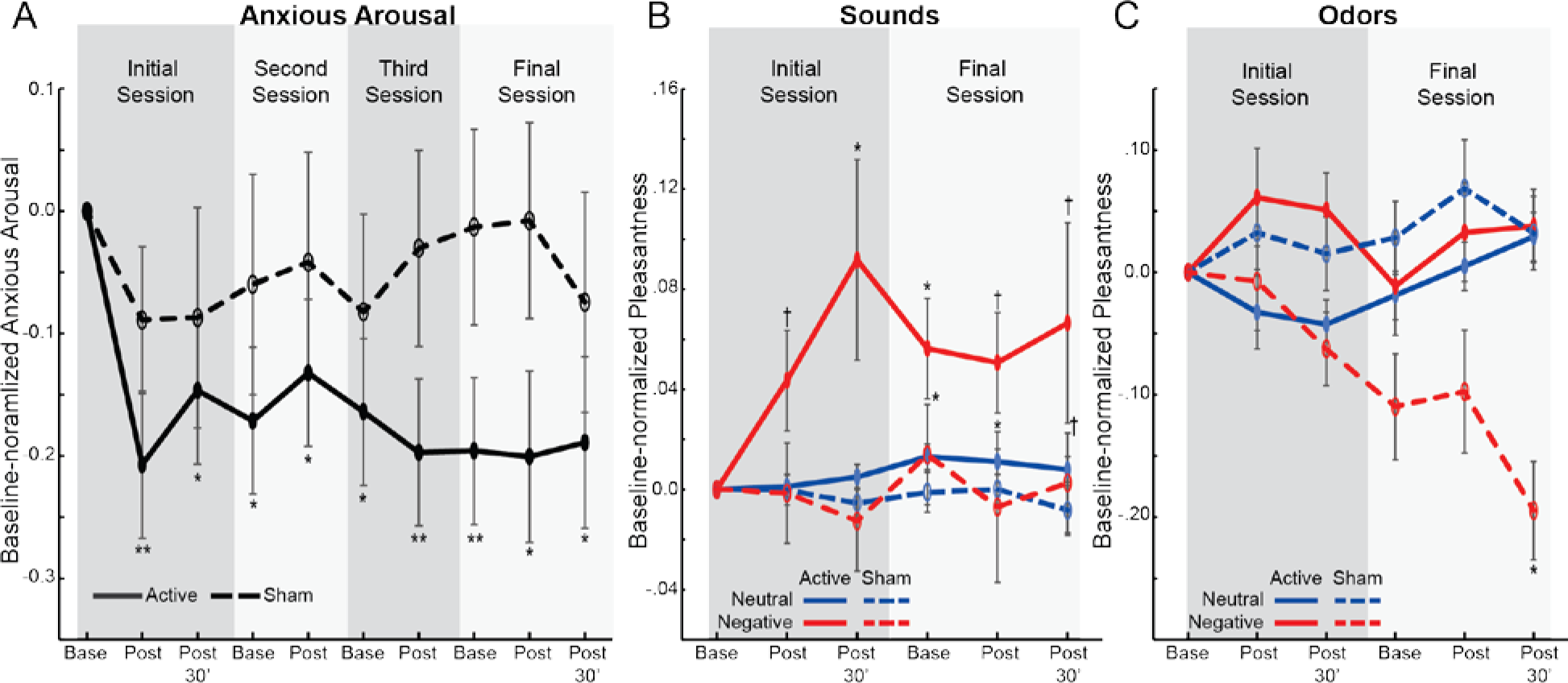
Clinical outcomes. A) Anxious arousal ratings, normalized to initial baseline ratings, for each group at each assessment over the four sessions/days. B-C) Pleasantness ratings of neutral and negative B) sounds and C) odors, normalized to initial baseline ratings. Error bars = SEM. \**p* < .05; \*\**p* < .01; † *p* < .1. All p’s are relative to the initial baseline. Note, the groups did not differ in the initial baseline ratings (*p’*s > .12).

### tACS increased perceived pleasantness of sensory stimuli

#### Auditory Stimuli

An omnibus rANOVA (Valence, Intensity, Session, Time, and Group) on pleasantness ratings of auditory stimuli revealed a main effect of Group (Active > Sham; *F*_*1, 32*_ = 4.19, *p* = .049, *η*_p_^2^ = .12), which was further characterized by a Session × Time × Group interaction (*F*_*1.57, 50.17*_ = 3.84, *p* = .038, *η*_p_^2^ = .11; Figure 3B). Breaking down the interaction by Session, an interaction of Time × Group emerged in the initial (*F*_*1.70, 54.28*_ = 4.10, *p* = .028, *η*_p_^2^ = .11) but not final (*p* = .707) session. That is, following the initial tACS (but not sham stimulation, *p* = .644), auditory stimuli were perceived as more pleasant (*F*_*1.35, 21.59*_ = 4.10, *p* = .045, *η*_p_^2^ = .20), immediately (*p* = .073) and 30 minutes (*p* = .040) after stimulation. Although no additional increases were seen after the final stimulation, final baseline ratings were significantly higher than initial baseline ratings, *t* = −2.43, *p* = .027, and ratings remained elevated post-stimulation (immediately, *p* = .053; 30 minutes,p = .083), suggesting sustained increases in pleasantness at the final session. No such between- or within-session changes were seen in the Sham group (*p’*s > .572). Therefore, sound pleasantness ratings paralleled anxious arousal ratings, increasing after the initial tACS and persisting through the last assessment on Day 4.

#### Olfactory Stimuli

A similar omnibus rANOVA on pleasantness ratings of olfactory stimuli revealed an interaction of Valence × Group (*F*_*1, 32*_ = 8.51, *p* = .006, *η*_p_^2^ = .21), which was further qualified by a Valence × Time × Group interaction (*F*_*1.75, 55.95*_ = 5.20, *p* = .011, *η*_p_^2^ = .14; Figure 3C). Follow-up ANOVAs revealed a Time × Group interaction for negative (*F*_*1.82, 58.10*_ = 4.60, *p* = .017, *η*_p_^2^ = .13) but not neutral (*p* = .230) odor ratings. That is, the Active group showed increases in pleasantness of negative odors (*F*_*1.88, 30.14*_ = 3.15, *p* = .060, *η*_p_^2^ = .16), both immediately (*p* = .050) and 30 minutes (*p* = .066) after stimulation. In contrast, the Sham group showed an opposing decrease in pleasantness of negative odors (*F*_*1.72, 27.56*_ = 3.32, *p* = .058, *η*_p_^2^ = .17). There were no Session × Group interactions or baseline shifts (*p’*s > .202) to indicate long-term, between-session effects.

## DISCUSSION

Administering repeated α-tACS over four days, we demonstrated both immediate and lasting effects of tACS. posterior→frontal alpha-frequency connectivity (in the right hemisphere) increased in the active (vs. sham control) group immediately and 30 minutes post-stimulation, which, indicative of a long-term effect of tACS, persisted at least 24 hours, resulting in a shifted baseline of posterior→frontal alpha influence. In parallel, anxious arousal decreased and perceived pleasantness of auditory stimuli increased immediately, 30 minutes, and 24 hours after α-tACS. In contrast, local occipito-parietal alpha power showed transient (immediately and 30 minutes post-stimulation) increases only, as did pleasantness ratings of negative olfactory stimuli. Therefore, the present study identifies long-term (> 24 hours) neural plasticity and clinically-relevant emotional improvement by tACS, providing strong support for its application in clinical interventions. In addition, the temporal dissociation between alpha power and connectivity implicates disparate mechanisms underlying tACS effects on neural oscillations, calling for further attention to oscillatory networks in response to tACS.

The acute (immediate and 30 minutes after) enhancement of local (occipito-parietal) alpha power was evident in both Sessions 1 and 4, corroborating previous reports of acute responses of posterior alpha oscillations to α-tACS (13, 17, 18) while highlighting the robustness of such effects. Interestingly, bin-by-bin examination of the tACS effect across the entire frequency spectrum (1-50 Hz) selectively isolated power enhancement in the lower half of the alpha range (8-10.5 Hz; Figure 2A), consistent with previous reports of left-ward shifting of alpha peaks by α-tACS (19) and a general trend of high alpha powers being associated with lower alpha peaks (46). Nevertheless, despite the repeated stimulation over consecutive days, we failed to observe long-term (> 24 hours) effects on local alpha power. It is possible that the current dosage was still at the lower end of the dose-response curve such that a dose-effect explanation may still account for the lack of lasting effects. However, the long-term effects in alpha connectivity seem to refute this account, indicating that the dosage was sufficient for generating lasting neural plasticity. Therefore, the shorted-lived alpha power increase likely reflects processes related to neural entrainment, which transiently boosted alpha power by temporally synchronizing occipito-parietal cortical oscillations to the exogenous stimulation frequency (20, 25). As such, at the cessation of tACS, the thalamo-cortical network, a unitary generator for alpha oscillations via dense bidirectional connections (28, 29), would reverse such local cortical entrainment via driving inputs from deep thalamic oscillators, resetting cortical alpha activity to the endogenous level.

It is known that alpha oscillations dominate the awake restful state and play critical roles in various neural and mental activities (38, 47–51). Conversely, aberrant resting-state alpha activity has been observed in multiple neuropsychiatric disorders (10, 11). In this sense, the resistance of cortical alpha oscillators to neural plastic changes due to tACS accentuates the stability of baseline alpha oscillations as well as the resilient, self-regulating internal network of alpha generators in healthy individuals (28, 29, 52, 53). Such qualities implicate a conserved, closed-loop (potentially encapsulated) system of alpha oscillators, which would serve an essential function in preserving neural homeostasis and mental regularity (54).

In contrast to the stable local alpha power, posterior→frontal GC connectivity in the alpha frequency exhibited both immediate and lasting changes, representing some of the first empirical evidence for long-term plasticity induced by tACS. The immediate effects replicated a recent study showing an immediate increase in resting-state connectivity between parietal and frontal cortices following α-tACS (32). Alpha power and alpha oscillatory connectivity are often correlated (11), which was also confirmed in the current study. However, the dissociation between alpha power and connectivity changes suggests distinct mechanisms were at play. Specifically, tACS did not induce lasting alpha power enhancement, ruling out the possibility that augmented alpha connectivity was the result of a stronger posterior sender (i.e., the occipitoparietal cortex). Rather, the evidence favored the interpretation that tACS improved the efficiency of alpha oscillatory propagation across long-range networks such that with alpha power at the sender remaining constant, directed transmission of alpha oscillations to the distant receiver was intensified.

This improved posterior→frontal alpha transmission efficiency is likely mediated by synaptic plasticity induced by tACS, via mechanisms such as STDP (15, 18, 21, 22, 55), that is known to be conducive to enduring Hebbian synaptic plasticity and lasting increases in cortico-cortical oscillatory interactions (30, 56). Given that such cortico-cortical connectivity operates outside the relatively encapsulated thalamo-cortical loop that constrains local cortical oscillations, its plasticity can outlast the effect of local alpha entrainment. This account concurs with the notion that tACS modulates electric fields in the cortex and thus directly modifies feedforward and feedback interactions across neurons to the extent that the dynamics of macroscopic, global networks are altered (15, 55). Given the role of alpha oscillations in long-range inter-regional interactions, augmented alpha rhythmic electric fields by α-tACS in the occipito-parietal cortex would generate pronounced connectivity plasticity in long-range pathways. In particular, the posterior→frontal alpha oscillatory causal connectivity exhibited maximal gain, akin to its dominance at resting long-range interactions. In contrast, connectivity in the opposite direction (frontal→posterior) showed minimal changes by tACS (see Supplemental materials).

Paralleling the alpha GC connectivity increases, anxious arousal reduced both immediately and 24 hours after tACS. Notably, we assessed anxious arousal every day in the four-day period and observed sustained reductions in anxious arousal over the days, highlighting a reliable and robust clinical effect. Evidently, these clinical effects outlasted local alpha power enhancement and so were likely mediated by strengthened posterior→frontal alpha connectivity. Long-range alpha oscillatory projections are thought to be closely associated with two major resting-state networks, the default mode network (DMN) and salience network (SN), which play critical roles in emotion processing and arousal response (57, 58) and whose activation levels are positively correlated with the strength of alpha oscillations (4, 39, 59–63). Therefore, tACS may modulate anxious arousal by facilitating the activity of these networks.

Similarly, we observed immediate and lasting increases in perceived pleasantness of sounds, especially negative sounds, in the tACS group (Figure 3B). Alpha oscillations are instrumental in regulating sensory processing and perception (31, 47, 64, 65) such that altered alpha activity via α-tACS could modify sensory analysis of these stimuli. In addition, auditory processing is widely known to be sensitive to states of arousal and alertness, as demonstrated by robust startle responses (41, 66) and arousal-related sensory gating (42). Therefore, reduced arousal states through tACS, potentially via augmentation of the DMN and SN and connectivity between visual and auditory cortices, can tone down aversive responses to auditory stimuli.

Interestingly, perceived pleasantness of negative odors increased only transiently, similar to the temporal profile of alpha power enhancement. Unlike the auditory system that is neocortical and has direct and close connections with the visual cortex (the current tACS target) (67–69), the olfactory system is largely subcortical or paleocortical without direct connections with the visual cortex (43–45). Given the lack of intrinsic connections between olfactory and visual structures, tACS-induced faciliation of cortico-cortical connecitivity may not directly impact olfactory nerual activity, thereby limiting the gain in olfactory affective perception. Rather, only short-living effects emerged, predominantly in negative odors, reflecting brief adaptation presumably mediated by transient effects of alpha enhancement.

Increasing recognition of aberrant oscillatory dynamics in the pathology of various neuropsychiatric disorders has spawned strong interests in directly modulating the endogenous oscillations within these “oscillopathies” (10, 70, 71). Furthermore, network views of psychopathology have gained rapid traction in recent years, advancing theorization of mental disorders from local neural aberrations to large-scale circuits anomalies (11, 72, 73). In fact, while we recently uncovered both deficient posterior alpha power and posterior→frontal alpha causal connectivity in posttraumatic stress disorder (PTSD), only the latter was directly correlated with the level of heightened anxious arousal and sensory hypervigilance that characterize PTSD (11). While local alpha power may be inherently stable and thus resistant to tACS modulation, lasting plasticity in long-range connectivity via tACS would effectively target network-related neuropathophysiology, promising a novel, transdiagnostic line of neuromodulatory treatment for neurological and psychiatric symptomatology rooted in aberrant oscillatory networks. In this regard, it is sensible to call for attention and interest to effects on oscillatory connectivity in future tACS research and application.

## METHODS

### Participants

38 healthy volunteers (18 female, 19.7 ± 2.0 years of age) participated in this study after providing written, informed consent approved by the Florida State University Institutional Review Board. No participants reported a history of neurological or psychiatric disorders and all were deemed tACS compatible (e.g. no metal implants, neurological surgery, or pregnancy). Participants were randomly assigned to two groups, an Active (*N* = 21) and a Sham control group (*N* = 17). Four Active-group participants did not complete the ratings or questionnaires, and were therefore excluded from the corresponding analyses. Groups did not differ in age or gender distribution (*p’*s > .14). Participants were blind to their group assignment.

### Ratings and questionnaires

#### Pleasantness ratings

Auditory and olfactory stimuli were rated for perceived pleasantness on a visual analog scale presented on a computer monitor (0-100 from most unpleasant to most pleasant). Auditory stimuli included a neutral (flat tone) and two negative (screaming and vomiting) sounds, delivered through headphones for one second each. Olfactory stimuli included a neutral (acetophenone, 5% l/l; diluted in mineral oil) and a negative (burning rubber; ScentAir™, Charlotte, NC) odor, delivered through a bottle. All stimuli were presented at three different intensities: weak, medium, and loud.

#### Subjective units of anxious arousal

Participants rated their current state of anxious arousal using a visual analog scale from 0 (not at all) to 100 (extremely), in accordance with standard ratings of Subjective Units of Distress (74).

### tACS

tACS was administered using a Soterix Medical 4x1 high-density (HD) transcranial electrical stimulation system (New York, NY, USA). Stimulation electrodes were placed in a 4x1 montage over occipito-parietal sites, with the central electrode receiving electrical current from the four surrounding electrodes (Figure 1A). Stimulation sites were selected to maximally target the dorsal extrastriate where deficient alpha oscillations in patients with posttraumatic stress disorder (PTSD) have been observed (11). Current flow modeling (Soterix Medical HDExplore Software, New York, USA) indicated maximal electric field intensity (0.21 V/m; with a 2 mA stimulation current) in the right dorsal occipital cortex (peak voxel: 29, −91, 18; Montreal Neurological Institute coordinates; Fig. 1A). Electric field intensity did not exceed 0.02 V/m in frontal regions.

Stimulation was administered for 30 minutes with a 2 mA sinusoidal current oscillating at individual participants’ baseline occipito-parietal peak alpha frequency (PAF). The PAF was identified as the peak frequency within the alpha-range (8-12 Hz) with a 0.5 Hz frequency resolution across occipito-parietal electrodes during the initial (session 1) baseline EEG recordings. Individual PAFs ranged from 8 to 11.5 Hz (*M* = 9.93, *SD* = 0.66 Hz) in the sample, equivalent between the two groups (*p* = .853). All participants were familiarized with tACS-induced skin sensations with a brief, 30 second pulse of random noise stimulation. During stimulation, the current was ramped up to 2 mA over a span of 10 seconds. Stimulation was then discontinued in Sham participants, and reintroduced during the last 10 seconds to mitigate awareness of experimental condition. Lastly, as tACS effects are especially pronounced in behavioral states with suppressed alpha power (23), participants were eyes-open, playing a simple computer game (Solitaire) throughout the procedure.

### Procedure

The experiment consisted of four sessions, each separated by 24 hours (Figure 1B). During the first and last sessions, participants started with a subjective rating of anxious arousal, followed by a two-minute resting, eyes-open EEG recording and then pleasantness ratings. Participant then underwent a tACS session, and then repeated a set of arousal ratings, resting EEG recordings, and pleasantness ratings. After a 30-minute break, participants again repeated a set of arousal ratings, resting EEG recordings, and pleasantness ratings. The second and third sessions included a rating of anxious arousal, a tACS session, and another rating of anxious arousal.

### EEG acquisition and analyses

EEG data were recorded from a 32-channel BrainProducts actiChamp system (1000 Hz sampling rate, 0.05 - 200 Hz online bandpass filter, referenced to the FCz channel). Electrooculogram (EOG) was recorded using four electrodes with vertical and horizontal bipolar derivations. EEG/EOG data were downsampled to 250 Hz, high-pass (1 Hz) and notch (60 Hz) filtered. We applied the *Fully Automated Statistical Thresholding for EEG artifact Rejection* (FASTER) algorithm for artifact detection and rejection (75).

#### Power analyses

EEG oscillation powers were computed for individual channels for each epoch (1-sec) using the multitaper spectral estimation technique (76). Alpha (8-12 Hz) powers were normalized by the mean power for the global spectrum (1-50 Hz) within each epoch. Alpha powers were extracted from occipito-parietal (right, middle, and left) electrodes, where they were maximally distributed (31, 47, 48).

### Directed alpha-frequency connectivity (Granger causality) analyses

Alpha-frequency Granger causality (GC) analysis (33, 34) was performed to assess posterior→frontal relay of alpha oscillatory activity. Following transformation into reference-free, current source density (CSD) data, scalp powers of ipsilateral frontal-posterior pairs were submitted to bivariate autoregressive (AR) modeling, from which Granger causality spectra were derived (34, 77).

### Statistical analyses

Occipito-parietal alpha power and feedforward alpha GC at each time point (*t*) were decibel-normalized relative to the initial (*i*), first-session pre-tACS, baseline values [power_*db*_ = 10*log10(power_*t*_/power_*i*_); GC_*db*_ = 10*log10(GC_*t*_/GC_*i*_) ] to control for variations in baseline activity (18, 19, 78). Note, the groups did not differ in initial baseline alpha power or GC (*p’*s > .19). The decibel-normalized values were submitted to omnibus rANOVAs of Site (power: left, middle, right; GC: left, right hemisphere), Session (initial, final), Time (pre, post, post-30 minutes), and Group (active, sham). Of particular relevance to our hypotheses were the interactions of Group with Time and/or Session, reflecting within-session (immediate and 30 minutes after) and between-session (accumulative and lasting) effects, respectively.

Pleasantness and anxious arousal ratings at each time point (*t*) were normalized using the difference-over-sum method to the initial (*i*) baseline [(rating_*t*_ ™ rating_*i*_)/(rating_*t*_ + rating_*i*_)] to account for variations in baseline ratings (79). The groups did not differ in initial baseline ratings (*p’*s > .12). Pleasantness ratings were submitted to omnibus rANOVAs of Valence (neutral, threat), Intensity (strong, medium, weak), Session (initial, final), Time (pre, post, post-30 minutes), and Group. Ratings of anxious arousal were submitted to an omnibus rANOVA of Session (initial, second, third, final), Time (pre, post) and Group. As ratings were acquired only before and immediately after tACS in the second and third sessions, we collapsed the immediate and 30-minute ratings for the initial and final sessions into a single post-stimulation rating. There was no difference between immediate and 30-minute ratings in these two sessions (*p’*s > .36). Individual differences in trait anxiety were removed by entering BIS scores as a covariate.

To isolate baseline shifts from the initial to the final session, thereby indicating long-term tACS effects (> 24 hours), we also performed ANOVAs of Session and Group on the prestimulation (baseline) data for all the dependent variables. Finally, multiple-comparison t-tests were controlled for Type I error using the false discovery rate/FDR criterion.

## Acknowledgement

This research was supported by the National Institute of Mental Health R01MH093413 (W.L.) and the FSU Chemical Senses Training (CTP) Grant Award T32DC000044 (K.C.) from the National Institutes of Health (NIH/NIDCD).

